# Receptor-specific interactome as a hub for rapid cue-induced selective translation in axons

**DOI:** 10.1101/673798

**Authors:** Max Koppers, Roberta Cagnetta, Toshiaki Shigeoka, Lucia C.S. Wunderlich, Sixian Zhao, Michael S. Minett, Anaïs Bellon, Clemens F. Kaminski, John. G. Flanagan, Christine E. Holt

## Abstract

During neuronal wiring, extrinsic cues trigger the local translation of specific mRNAs in axons via cell surface receptors. The coupling of ribosomes to receptors has been proposed as a mechanism linking signals to local translation but it is not known how broadly this mechanism operates, nor whether it can selectively regulate mRNA translation. We report that receptor-ribosome coupling is employed by multiple guidance cue receptors and this interaction is mRNA-dependent. We find that different receptors bind to distinct sets of mRNAs and RNA-binding proteins. Cue stimulation induces rapid dissociation of ribosomes from receptors and the selective translation of receptor-specific mRNAs in retinal axon growth cones. Further, we show that receptor-ribosome dissociation and cue-induced selective translation are inhibited by simultaneous exposure to translation-repressive cues, suggesting a novel mode of signal integration. Our findings reveal receptor-specific interactomes and provide a general model for the rapid, localized and selective control of cue-induced translation.

## Introduction

mRNA localization and local translation are major determinants of the local proteome (Zappulo et al., 2017). This seems particularly important for morphologically complex cells such as neurons, where the axonal sub-compartment and its tip, the growth cone, often far away from the cell body, can rapidly perform specialized functions (Holt and Schuman, 2013). During neuronal wiring, specific interactions between extrinsic cues and receptors mediate guidance of axons to their proper target area, where they also regulate axon branching (Stoeckli, 2018, Manitt et al., 2009, Marshak et al., 2007, Cioni et al., 2013). The rapid axonal responses to several guidance cues require local protein synthesis (Jung et al., 2012, Campbell and Holt, 2001). Attractive guidance cues, such as Netrin-1, trigger axonal translation of mRNAs encoding proteins that facilitate actin assembly whereas repulsive cues trigger the local synthesis of cytoskeletal proteins involved in actin disassembly (Leung et al., 2006, Wu et al., 2005, Piper et al., 2006). This cue-specific mode of translation enables growth cones to steer differentially towards/away from a polarised cue (Lin and Holt, 2007, Lin and Holt, 2008). Unbiased detection of newly synthesized proteins in the axon compartment has revealed further complexity showing that different guidance cues stimulate the regulation of distinct signature sets of >100 axonal nascent proteins within just 5 mins, many of which are not cytoskeletal-related (Leung et al., 2006, Yao et al., 2006, Wu et al., 2005, Cagnetta et al., 2018, Cioni et al., 2018). However, the mechanisms underlying the localization and selectivity of translation are unclear. Several mechanisms are known to control different aspects of axonal translation, including microRNA regulation (Bellon et al., 2017), mRNA modification (Yu et al., 2018), modulation of the phosphorylation of eukaryotic initiation factors (Cagnetta et al., 2019), RNA-binding protein (RBP) phosphorylation (Sasaki et al., 2010, Lepelletier et al., 2017, Huttelmaier et al., 2005) and receptor-ribosome coupling (Tcherkezian et al., 2010). The latter is a particularly direct and attractive mechanism to link cue-specific signalling to differential mRNA translation. However, this mechanism has been shown only for the Netrin-1 receptor, deleted in colorectal cancer (DCC), in commissural axon growth cones and HEK293 cells (Tcherkezian et al., 2010) and it is unknown whether receptor-ribosome coupling is a widespread mechanism used by different receptors and in different cell types, and whether it regulates selective local translation.

Here, we show in retinal ganglion cell (RGC) axon growth cones that receptor-ribosome coupling is used by several different axon guidance receptors (DCC, Neuropilin-1 and Robo2), indicative of a common mechanism. Upon stimulation by specific cues, ribosomes dissociate from their receptors within 2 minutes. Interestingly, the receptor-ribosome interaction is mRNA-dependent, and immunoprecipitation (IP) reveals that distinct receptors associate with specific RNA-binding proteins (RBPs) and subsets of mRNAs suggesting that this may indeed be the basis for the selective translation of receptor-specific mRNAs. Finally, we show that EphrinA1 co-stimulation blocks Netrin-1-induced DCC receptor-ribosome dissociation and Netrin-1-mediated selective translation in axons, providing a mechanism to integrate different signals. Together, this study provides evidence that receptor-ribosome coupling is a common mechanism for the rapid regulation of axonal translation downstream of different guidance cues and suggests receptor-specific interactomes act as a hub to regulate the localized and selective cue-induced mRNA translation.

## Results

### Multiple guidance cue receptors interact with ribosomes

In retinal axons, Netrin-1 and Sema3A mediate growth cone steering and branching (Campbell and Holt, 2001, Manitt et al., 2009, Campbell et al., 2001). Specifically, the rapid chemotropic responses to Netrin-1 and Sema3A are mediated, at least in part, by local translation (Campbell and Holt, 2001). The Netrin-1 receptor, DCC, was previously reported to associate with ribosomes in spinal commissural axon growth cones (Tcherkezian et al., 2010). We first asked whether the interaction of DCC with ribosomes is conserved in a different system and cell type, and explored the possibility that the Sema3A receptor, Neuropilin-1 (Nrp1), also interacts with ribosomes in this system. To do this, we performed immunoprecipitation (IP) of endogenous DCC and Nrp1 from *Xenopus laevis* embryonic brains and eyes followed by mass-spectrometry (LC-MS/MS) analysis of eluted samples. Each IP was performed in triplicate and after raw data processing using MaxQuant software, we determined statistically significant interactors of DCC and Nrp1 compared to an IgG control pulldown using label-free (LFQ) intensities and Perseus software analysis (Figure 1A). Gene-ontology (GO) enrichment analysis revealed that ‘structural constituent of ribosomes’ appeared as the most prominently enriched category in both DCC and Nrp1 pulldowns, indicating that both receptors can interact with ribosomal proteins (Figure 1B). Specifically, 75 out of 79 ribosomal proteins (94.9%) were detected in the DCC and Nrp1 pulldowns. Of these, 51 and 33 RPs were identified as statistically enriched interactors for Nrp1 and DCC, respectively, compared to IgG control pulldowns. There was no bias towards small or large ribosomal subunit proteins (Figure 1A, red dots). The GO analysis also revealed the presence of other groups shared between the receptors, such as ‘vesicle-mediated transport’ (Figure 1B). Interestingly, some categories of proteins were enriched for only one of the receptors, for example the ‘phosphoprotein phosphatase activity’ GO term was significantly enriched only in the DCC pulldown and the ‘barbed-end actin filament capping’ GO term was enriched only in the Nrp1 pulldown (Figure 1B). To confirm the interaction between receptors and ribosomal proteins, we performed Western blot (WB) after IP and validated that both DCC and Nrp1 interact with small (40S) and large (60S) ribosomal subunit proteins (Figure 1C-D). These interactions appear to be conserved, as endogenous IP from the human neuronal cell line SH-SY5Y, which expresses both DCC and Nrp1, also shows ribosomal protein co-precipitation after pulldown of the endogenous receptor (Figure S1A).

**Figure 1.**
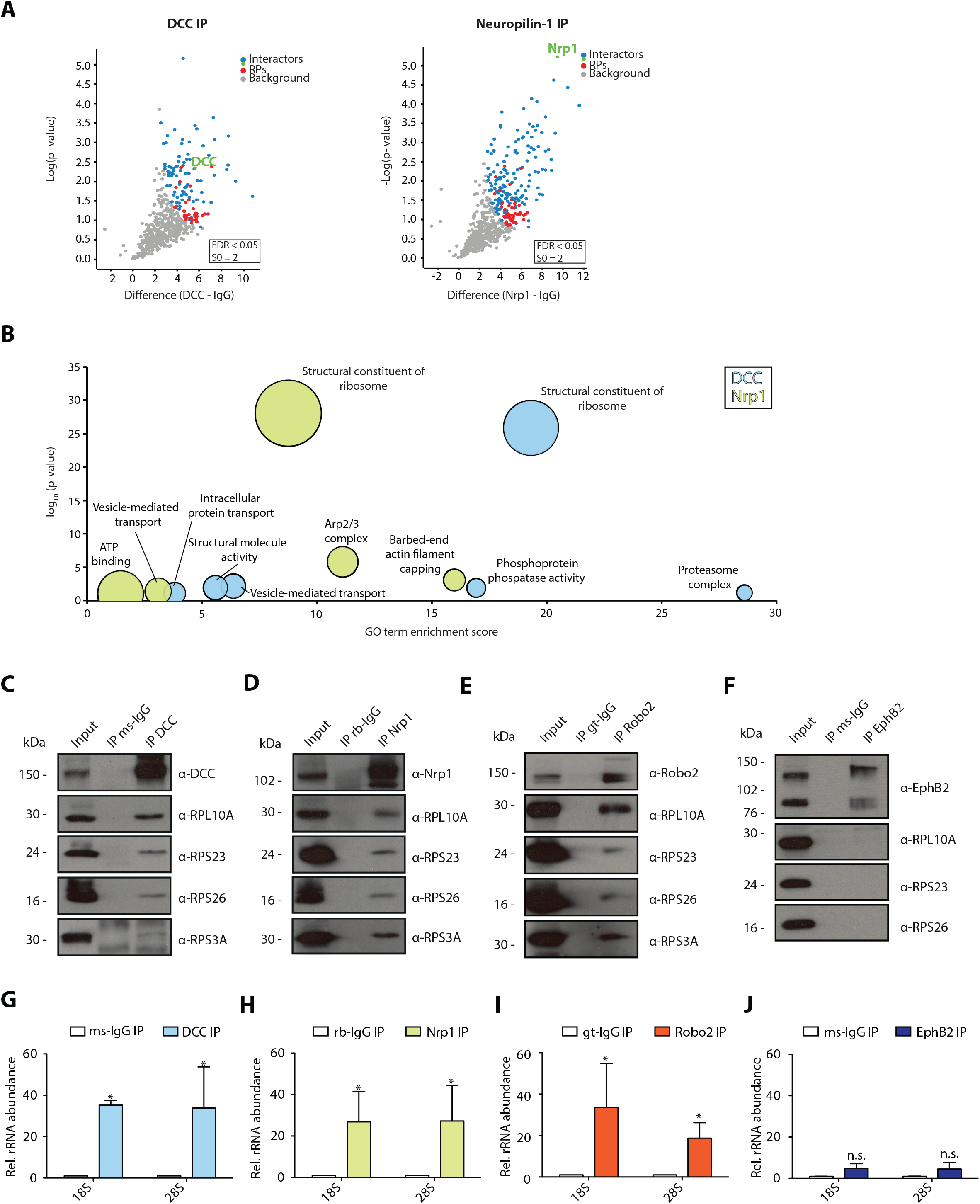
Multiple guidance cue receptors interact with ribosomes. (A) Volcano plots showing statistically enriched proteins in DCC-IP and Nrp1-IP samples identified by permutation-based FDR-corrected t-test based on three biological replicates. The LFQ intensity of the DCC or Nrp1 pulldowns over IgG pulldowns are plotted against the - log10 p-value. FDR <0.05; S0 = 2. (B) Gene enrichment analysis of statistically enriched proteins in the DCC and Nrp1 pulldown samples. (C-F) Western blot validation of RP co-immunoprecipitation with DCC, Nrp1 and Robo2 but not with EphB2. (G-J) Relative 18S and 28S ribosomal RNA abundance after control (IgG) pulldown or receptors pulldowns shows enrichment of rRNA in DCC, Nrp1, and Robo2 but not EphB2 pulldowns (unpaired two-tailed t-test; three biological replicates). Error bars indicate standard deviation; * p<0.05.

In addition to DCC and Nrp1, Roundabout 2 (Robo2) triggers local protein synthesis after binding to the guidance cue Slit2 (Piper et al., 2006). Therefore, we asked whether Robo2 also interacts with ribosomal proteins. WB after IP from *Xenopus* embryonic brains and eyes showed that Robo2 also interacts with ribosomal proteins of both subunits (Figure 1E). Growth cone collapse via EphB2 is not mediated by local protein synthesis (Mann et al., 2003), raising the possibility that only receptors that require local protein synthesis for their action on growth cones are coupled to ribosomes. In support of this idea, we could not detect co-IP of ribosomal proteins with EphB2 in *Xenopus* embryonic brains and eyes, indicating that not all guidance receptors interact with ribosomal proteins (Figure 1F).

To confirm that receptors bind to ribosomes or ribosomal subunits and not free ribosomal proteins, we isolated RNA after IP and performed quantitative-RT-PCR (qPCR) for 18S (40S small ribosomal subunit) and 28S (60S large ribosomal subunit) ribosomal RNA (rRNA), which should be present only in intact ribosomal subunits in the cytoplasm. Consistent with the WB results, DCC, Nrp1 and Robo2, but not EphB2, exhibit a significant fold enrichment of both 18S rRNA and 28S rRNA compared to an IgG control pulldown in both *Xenopus* brains (Figure 1G-J) and SH-SY5Y cells (Figure S1B). Collectively, these findings reveal that multiple receptors known to trigger local protein synthesis can associate with ribosomes.

### Possible heterogeneity between DCC- and Nrp1-bound ribosomes

Proteomic studies in yeast and mammalian cells have shown evidence for ‘specialized’ ribosomes, which exhibit different stoichiometries for certain ribosomal proteins and are thereby tuned to preferentially translate specific subsets of mRNAs (Shi et al., 2017, Segev and Gerst, 2018, Ferretti et al., 2017). Therefore, an intriguing hypothesis for developing neurons is that each guidance cue receptor might act as a trans-factor binding to ribosomes of different compositions, which regulate selective translation. To preliminarily test this hypothesis, we used MS-based label-free quantification, which has been previously used to determine stoichiometries of individual proteins in protein complexes, and calculated the relative iBAQ values of all the ribosomal proteins detected (Schwanhausser et al., 2011, Smits et al., 2013). The analysis identified several candidate ribosomal proteins showing a different stoichiometry between the DCC- and Nrp1-bound ribosomes (Figure S1C-D). Of note, RPL4/uL4 is significantly more abundant in Nrp1 versus DCC pulldowns, and RPL39/eL39 is absent from Nrp1 pulldowns but present in 2 out of 3 DCC pulldown replicates (Figure S1C-D). These data suggest the possibility that ribosomes bound to DCC might have a different composition compared to those bound to Nrp1.

### Guidance cue receptors associate with ribosomes in a mRNA-dependent manner

To gain mechanistic insights into the interaction between receptors and ribosomes, we examined the co-sedimentation profiles of DCC and Nrp1 in *Xenopus* embryonic brains and eyes after sucrose gradient purification of ribosomes. Consistent with previous findings (Tcherkezian et al., 2010), DCC was prominent in 40S, 60S and 80S fractions but not in polysomal fractions (Figure S2A). Nrp1 was also found in 40S, 60S and 80S fractions, as well as some detection in polysomal fractions (Figure S2A), suggesting a possibly different association mechanism between the two receptors, or a different translational status of receptor-bound ribosomes. Both DCC and Nrp1 were also present in ribosome-free fractions indicating that not all receptor molecules are associated with ribosomes (Figure S2A). EDTA treatment, which dissociates the monosomes/polysomes into separate ribosomal subunits, shifted both DCC and Nrp1 to lighter fractions, supporting a valid association with ribosomes (Figure S2B).

Next, we used qPCR to further investigate the association of ribosomes to receptors. When IP samples were treated with EDTA before elution, the enrichment of 18S and 28S rRNA after receptor pulldown was significantly decreased for both DCC and Nrp1 (Figure 2A). One possibility for this decrease is that DCC and Nrp1 interact mainly with 80S ribosomes. Another possibility is that the binding of ribosomes to receptors is mRNA-dependent. To test the latter hypothesis, we treated the receptor pulldown samples with RNase A/T1, which digests mRNAs and releases any factors bound to ribosomes via mRNA (Simsek et al., 2017). We have previously shown that the concentration of RNase A/T1 used largely preserves the integrity of ribosomes although we cannot formally exclude that it may still partially cleave rRNA (Shigeoka et al., 2018). We found a significant decrease in the co-precipitation of 18S and 28S rRNA with receptors (Figure 2A). Consistent with these results, Western blot analysis of IP samples treated with RNaseA/T1 or EDTA treatment confirms the decrease in ribosomal protein binding after RNaseA/T1 or EDTA treatments for both DCC and Nrp1 (Figure 2B, C). Together, these results suggest that the interaction of receptors with ribosomes is largely mediated through mRNA.

**Figure 2.**
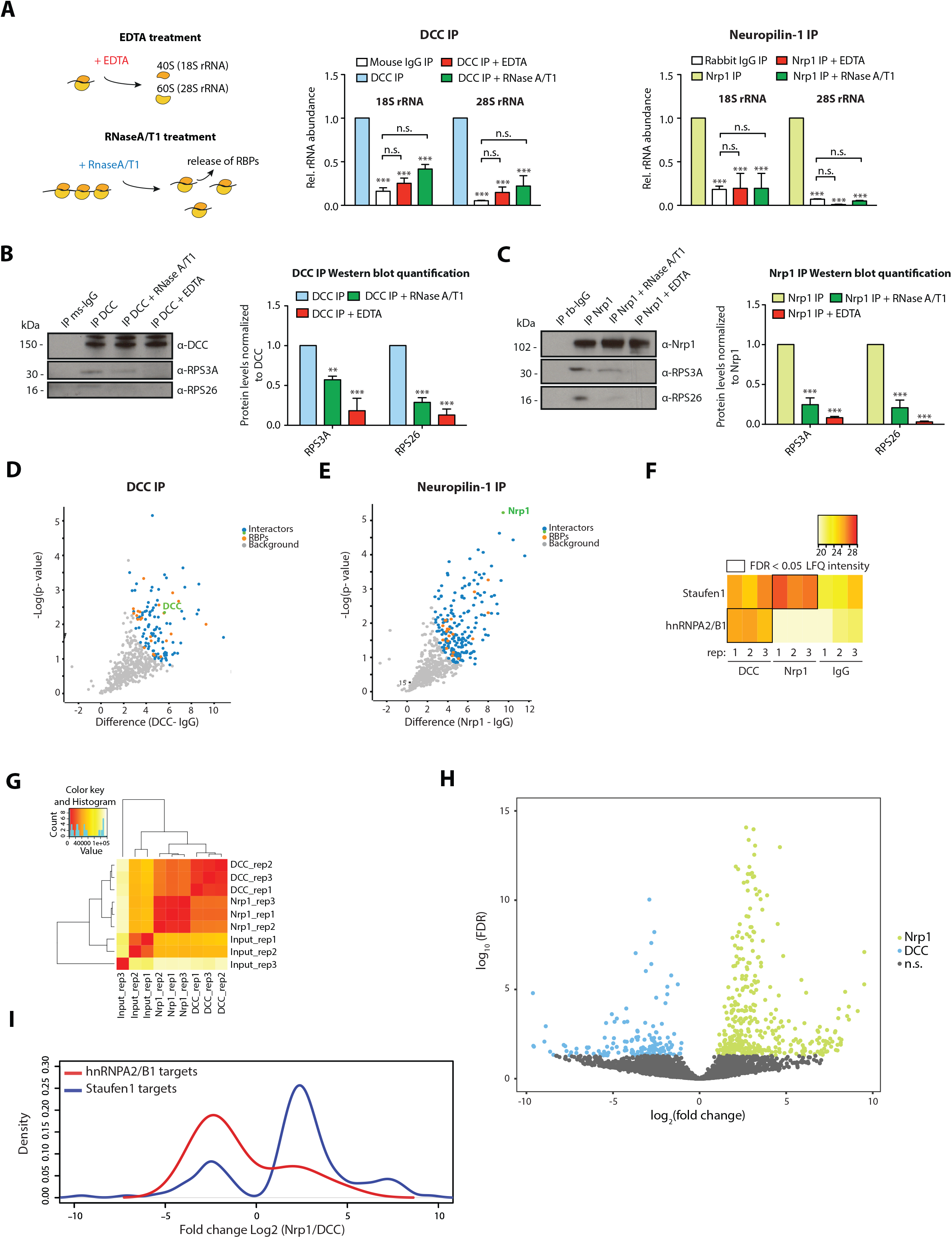
Receptor-ribosome coupling is mRNA dependent and DCC and Nrp1 bind to specific RBPs and mRNAs. (A) Relative 18S and 28S ribosomal RNA abundance after control (IgG) pulldown or receptors pulldowns with or without EDTA or RNase A/T1 treatments (two-way ANOVA with Bonferroni’s multiple comparisons test; three biological replicates; Error bars indicate standard deviation; ***p<0.0001). (B) Western blot analysis and quantification of ribosomal proteins after DCC and (C) Nrp1 pulldowns. (two-way ANOVA with Bonferroni’s multiple comparisons test; three biological replicates; Error bars indicate standard deviation; **p<0.01; ***p<0.0001). (D, E) Volcano plots indicating both DCC and Nrp1 bind significantly to different RBPs (orange dots). FDR <0.05; S0 = 2. (F) DCC and Nrp1 each bind to differentially to the RBPs Staufen1 and hnRNPA2B1. FDR <0.05. (G) Distance matrix showing a high correlation between replicates and a distinct signature between samples. (H) Volcano plot showing differential expression analysis for DCC and Nrp1 pulldowns. (I) Enrichment analysis plot of known RBP targets detected in RNA-sequencing data after DCC and Nrp1 pulldown.

### DCC and Nrp1 bind to specific RBPs and subsets of mRNAs

The mRNA-dependency of the receptor-ribosome interaction suggests that receptors may bind to ribosomes via mRNAs and/or RNA binding proteins. Indeed, our MS analysis revealed that several RBPs are significantly enriched after DCC or Nrp1 pulldown (Figure 2D, E, orange dots). Of 23 RBPs pulled down with DCC and 34 RBPs pulled down with Nrp1, only 10 are shared between the two receptors (Figure 2D, E). Several RBPs are significantly enriched in only one of the two receptor IPs. For example, Staufen1 is significantly enriched after Nrp1 IP, but not DCC IP (Figure 2F), whereas hnRNPA2/B1 is only detected after DCC IP, as confirmed by Western blot analysis (Figures 2F and S2C). Together with our finding that mRNA mediates the association of receptors with ribosomes, these results suggest a model in which receptors associate with specific RBPs, which bind specific mRNAs, and these mRNAs, in turn, recruit ribosomes.

Next, we examined if and which mRNAs can associate with DCC and Nrp1 by performing RNA-sequencing (RNA-seq) on RNAs isolated after DCC and Nrp1 IP. We used SH-SY5Y cells, a neuronal cell line, for these experiments to rule out that any detected differences in mRNAs are due to expression of DCC and Nrp1 in different cell types. Co-precipitation of RNA was observed in DCC and Nrp1 pulldowns but not in IgG control pulldowns (Figure S2D). A distance matrix analysis revealed that the experimental replicates clustered together for each receptor and we observed a distinct signature of detected mRNAs between DCC, Nrp1 or whole lysate input samples (Figure 2G). Differential expression analysis revealed that DCC and Nrp1 each differentially bind to specific subsets of mRNAs, with 541 mRNAs differentially binding between DCC and Nrp1 (158 mRNAs for DCC *vs* 383 mRNAs for Nrp1) (Figure 2H). GO enrichment analysis of differentially expressed mRNAs showed the enrichment of mRNAs involved in different processes for each receptor (Figure S2E, S2F).

Although these results rely on mRNA populations expressed in SH-SY5Y cells, we compared mRNAs that preferentially bind to DCC or Nrp1 (Figure 2H) with known mRNA targets of several RBPs (Staufen1, hnRNPA2B1, Elavl1 and Fxr1), which were identified by previous CLIP studies in other systems (Lebedeva et al., 2011, Martinez et al., 2016, Sugimoto et al., 2015, Ascano et al., 2012). We focused particularly on Staufen1 and hnRNPA2/B1 because our proteomic analysis revealed that Staufen1 is enriched after Nrp1 pulldown compared to DCC pulldown and hnRNPA2/B1 was only detected after DCC pulldown (Figure 2F). Our analysis revealed significant enrichment of known Staufen1 and hnRNPA2/B1 targets, respectively, in Nrp1 *versus* DCC pulldown, respectively (Figure 2I). Indeed, 41.1% percent of the significantly enriched DCC-precipitated RNAs and 43.1% percent of the significantly enriched Nrp1-precipitated mRNAs can be explained by known targets of four RBPs tested (Staufen1, hnRNPA2/B1, Elavl1 and Fxr1). Collectively, the results support a model where receptor-specific RBPs mediate differential association of mRNAs to receptors.

### Receptor-ribosome coupling occurs in RGC axonal growth cones and is cue-sensitive

A proposed key advantage of receptor-ribosome coupling is the precise spatiotemporal control of translation in subcellular compartments, such as axonal growth cones or discrete filopodia/branches (Tcherkezian et al., 2010). As our IP experiments were performed in whole brain lysates, we asked whether these interactions occur in retinal growth cones. To address this question, we cultured eye primordia from *Xenopus* embryos and performed immunocytochemistry and expansion microscopy (Chen et al., 2015) on retinal axons using antibodies against the intracellular domain of DCC and a ribosomal protein (Figure 3A). DCC and RPL5/uL18 partially colocalized in retinal growth cones and filopodia (Figure 3A, white arrowheads). Similarly, RPS3A/eS1 with Nrp1 co-localized in retinal growth cones (Figure 3B, white arrowheads). To further confirm receptor-ribosome coupling in axonal growth cones, we employed the proximity-ligation assay (PLA) (Soderberg et al., 2006), modified for use on retinal axons (Yoon et al., 2012), which reports the spatial coincidence (< 40nm) of two proteins of interest by using the respective antibodies. As a negative control, PLA was performed using DCC antibody with an IgG control antibody, which generated a very low amount of background PLA signal (Figure 3C, S3A). In contrast, we detected a strong PLA signal between DCC and RPL5/uL18, in line with previous findings (Konopacki et al., 2016), as well as with RPS4X/eS4 or RPL10A/uL1 (Figure 3C, S3A). Similarly, Nrp1 generated a strong PLA signal together with RPS3A/eS1 or RPS23/uS12, and no detectable PLA signal in the negative control (Nrp1-IgG) (Figure 3D). These results are consistent with the association of DCC and Nrp1 with ribosomal proteins. Given that EphB2 IP in *Xenopus* brain and eyes does not show any interaction with ribosomal proteins (Figure 1F), we tested whether this is conserved in retinal growth cones. Consistent with the IP results (Figure 1F) and with the EphB2-induced local protein synthesis independent growth cone collapse (Mann et al., 2003), PLA between EphB2 and RPL5/uL18 shows almost no detectable signal compared to DCC-RPL5/uL18 or Nrp1-RPS3A/eS1 in growth cones (Figure 3E).

**Figure 3.**
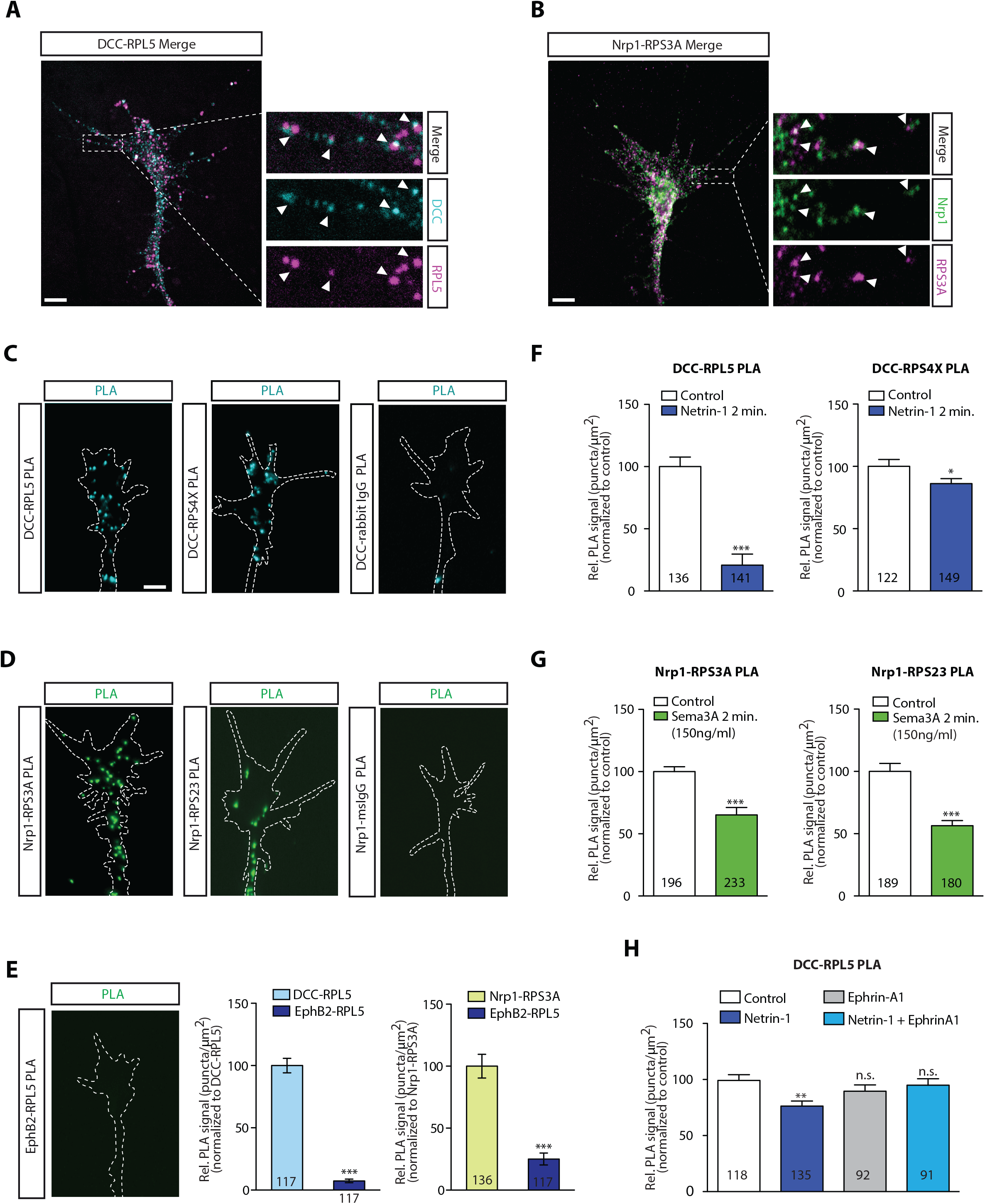
DCC and Nrp1 are in close proximity to ribosomes in axonal growth cones in a cue-dependent manner. (A) Expansion imaging shows partial colocalization of DCC and (B) Nrp1 with ribosomal proteins (Scale bars, 5 μm). (C) Representative proximity ligation assay signal in axonal growth cones between DCC and RPL5/uL18, RPS4X/eS4 or IgG control (Scale bars, 5 μm). (D) Representative proximity ligation assay signal in axonal growth cones between Nrp1 and RPS3A/eS1, RPS23/uS12 or IgG control (Scale bars, 5 μm). (E) EphB2 and RPL5/uL18 show a significantly lower amount of PLA signal in axonal growth cones compared to DCC-RPL5/uL18 or Nrp1-RPS23/uS12 (Mann-Whitney test; ***p<0.0001; Scale bars, 5 μm). (F, G) Quantification of PLA signal in cue-stimulated axonal growth cones relative to control (unpaired two-tailed t-test; error bars indicate SEM; ***p<0.0001; *p = 0.0423). (H) Relative PLA quantification of DCC and RPL5/uL18 compared to control after Netrin-1, EphrinA1, or co-stimulation (one-way ANOVA with Bonferroni’s multiple comparisons test; error bars indicate SEM; **p = 0.0058). For all PLA experiments, numbers in bars indicate amount of growth cones quantified collected from at least three independent experiments.

Previous work has shown that stimulation with the guidance cues Netrin-1 and Sema3A that bind DCC and Nrp1, respectively, triggers the remodelling of the nascent axonal proteome within 5 minutes (Cagnetta et al., 2018). Therefore, we investigated if the association between receptors and ribosomal proteins is cue-sensitive. Remarkably, the PLA signal between DCC and the ribosomal proteins RPL5/uL18 and RPS4X/eS4 decreased significantly in retinal axon growth cones after 2 min of Netrin-1 of stimulation (Figure 3F), suggesting a rapid dissociation of ribosomes from the receptor. It should be noted that DCC protein level does not change in response to 5 min Netrin-1 stimulation, whereas both RPL5/uL18 and RPS4X/eS4 are up-regulated in response to 5 min Netrin-1 stimulation (Cagnetta et al., 2018). Therefore, the decrease in the PLA signal in response to Netrin-1 may be underestimated. Similarly, the PLA signal between Nrp1 and RPS3A/eS1 and RPS23/uS12 decreases within 2 min stimulation with Sema3A (Figure 3G), at a concentration known to affect local axonal translation (Manns et al., 2012, Nedelec et al., 2012). The detected decrease in PLA signal should not be affected by Nrp1, RPS3A/eS1 and RPS23/uS12 protein levels as they are unchanged in response to 5 min Sema3A stimulation (Cagnetta et al., 2018). Higher concentration of Sema3A (700ng/ml) is known to induce protein synthesis-independent growth collapse (Nedelec et al., 2012, Manns et al., 2012) and puromycylation of newly synthesized proteins (Schmidt et al., 2009) in the presence of 700ng/ml Sema3A shows no increase in global translation in growth cones (Figure S3B), in contrast to stimulation with 150ng/ml Sema3A, which triggers 30% increase in local protein synthesis (Cagnetta et al., 2019). Interestingly, in line with this finding, stimulation with 700ng/ml Sema3A does not cause a rapid decrease in the Nrp1-RPS3A/eS1 PLA signal (Figure S3C), whereas stimulation with 150ng/ml Sema3A trigger ∼35% decrease in the Nrp1-RPS3A/eS1 PLA signal (Figure 3G).

During axon pathfinding and branching, axons encounter and integrate multiple cues, such as EphrinB2 and Netrin-1, known to generate a complex between the respective receptors (Morales and Kania, 2017, Dudanova and Klein, 2013, Poliak et al., 2015). The cue EphrinA1 has been reported to decrease local translation in hippocampal axons (Nie et al., 2010) and the rapid local translation of the Translationally controlled tumor protein (Tctp), which is up-regulated by Netrin-1 (Gouveia Roque and Holt, 2018). Therefore, we asked whether co-stimulation of EphrinA1 with Netrin-1 alters the dissociation of ribosomes from the Netrin-1 receptor, DCC. To address this question, we co-stimulated retinal axons with Netrin-1 and EphrinA1 and examined receptor-ribosome coupling by using the PLA approach. Whereas Netrin-1 induces a decrease in the DCC-RPL5/uL18 PLA signal within 2 min, both Ephrin-A1 stimulation alone and co-stimulation of Netrin-1 with EphrinA1 do not decrease the DCC-RPL5/uL18 PLA signal, indicating that the Netrin-1-induced dissociation of ribosomes from DCC is blocked by EphrinA1 co-stimulation (Figure 3H). By contrast, EphrinA1 co-stimulation with Sema3A does not block the Sema3A-induced decrease in the Nrp1-RPS23/uS12 PLA signal (Figure S3D).

Together, these findings extend receptor-ribosome coupling to retinal axon growth cones downstream of multiple receptors mediating local protein synthesis and point towards a model where dissociation of ribosomes from receptors is a shared mechanism among cue-induced local translation-dependent responses. Furthermore, the results reveal that integration of guidance cues can alter the receptor-ribosome dissociation, possibly by structural changes of the interacting receptors (Morales and Kania, 2017, Dudanova and Klein, 2013, Poliak et al., 2015).

### Integration of multiple cues can affect cue-induced selective translation of receptor-specific mRNAs

Our data showing EphrinA1 blocks the Netrin-1-induced ribosome dissociation from DCC, suggests that EphrinA1 may inhibit (selective) local translation induced by Netrin-1. To test this hypothesis, we examined the effect of cue integration of Netrin-1 and EphrinA1 on both global and selective local translation in growth cones. Axon-only cultures, after soma removal, were pulse-labeled with a low concentration of puromycin, a structural tRNA analogue that is incorporated into the C-terminus of nascent polypeptide chains. Subsequently, global translation was visualized after incubating with an anti-puromycin antibody and measured by quantitative immunofluorescence (Schmidt et al., 2009). In the culture conditions used in this study (Hopker et al., 1999), both Netrin-1 and EphrinA1 decrease global local translation in axons (Figure 4A-B). Consistent with this result, both cues decrease pERK1/2 levels (Figure S4A), an upstream activator of the TOR signalling pathway, which is known to regulate axonal protein synthesis (Campbell and Holt, 2003).

**Figure 4.**
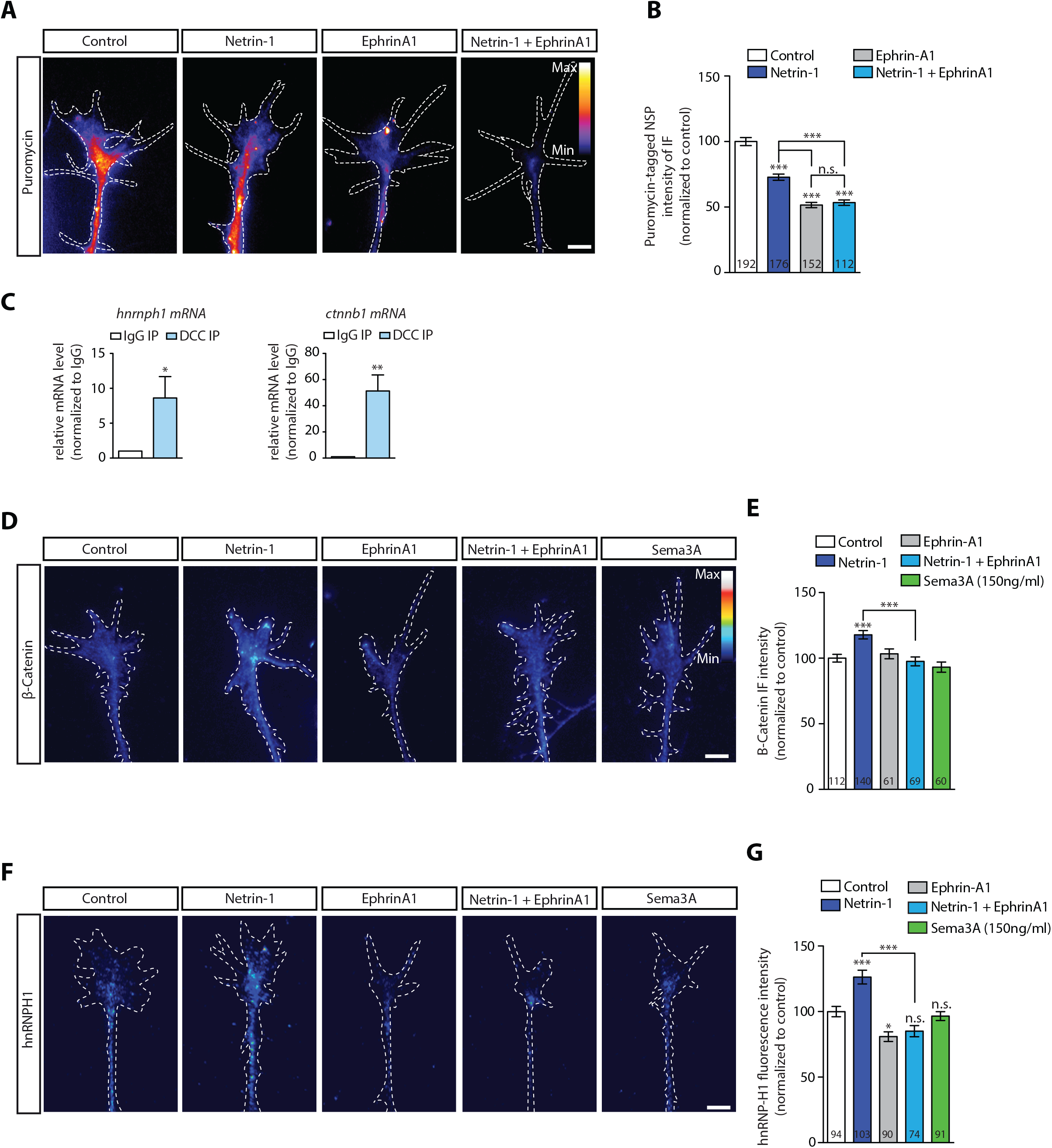
EphrinA1 co-stimulation blocks Netrin-1 induced receptor-ribosome dissociation and selective translation. (A, B) Puromycin QIF relative to control after Netrin-1, EphrinA1 or co-stimulation (one-way ANOVA with Bonferroni’s multiple comparisons test; error bars indicate SEM; ***p<0.0001). (C) Relative mRNA quantification after DCC IP of *hnrnph1* and *ctnnb1* mRNA (unpaired t-test with Welch’s corrections on dCT values; three biological replicates; error bars indicate SEM; *p=0.02 for *hnrnph1*; **p=0.0018 for *ctnnb1*). (D, E) B-Catenin QIF relative to control after Netrin-1, EphrinA1, Sema3A or Netrin-1 and EphrinA1 co-stimulation (one-way ANOVA with Bonferroni’s multiple comparisons test; error bars indicate SEM; ***p<0.0001). (F, G) hnRNPH1 QIF relative to control after Netrin-1, EphrinA1, Sema3A or Netrin-1 and EphrinA1 co-stimulation (one-way ANOVA with Bonferroni’s multiple comparisons test; error bars indicate SEM; ***p<0.0001; *p=0.0164). Scale bars, 5 μm. For all QIF experiments, numbers in bars indicate amount of growth cones quantified collected from at least three independent experiments.

Despite the decrease in global local translation, previous work has revealed that Netrin-1 can induce the rapid selective translation of specific mRNAs (Cagnetta et al., 2018, Shigeoka et al., 2018). The IP-RNA-seq data in SH-SY5Y cells had revealed that DCC associates with mRNAs encoding β-catenin (*ctnnb1*) and hnRNPH1 (*hnrnph1*) significantly more than with Nrp1. *Ctnnb1* and *hnrnph1* mRNAs have been detected in RGC axons by RNA-seq (Shigeoka et al., 2018). We next tested whether these mRNAs associate with DCC also in *Xenopus* brain and eyes, by carrying out IP followed by qPCR. The results showed significant enrichment of *ctnnb1* and *hnrnph1* mRNAs in DCC pulldown compared to an IgG pulldown, thus confirming their association with DCC (Figure 4C). Importantly, both β-catenin and hnRNPH1 proteins are selectively synthesised in response to 5 min Netrin-1 stimulation, but not Sema3A, as revealed by quantification of immunofluorescence (Figure 4D-G).

Similar to β-catenin and hnRNPH1, RPS14/uS11 is up-regulated in response to 5 min Netrin-1 stimulation, but not Sema3A (Cagnetta et al., 2018), as confirmed by quantification of immunofluorescence (Figure S4D). However, *rps14* mRNA was not detected to be associated with DCC in SH-SY5Y. Therefore, we asked whether this is due to interspecies differences (SH-SY5Y vs. *Xenopus*), or whether *rps14* is selectively translated via a DCC interactome-independent mechanism. To address this question, we carried out IP followed by qPCR in *Xenopus* brain and eyes, which confirmed *rps14* association to DCC (Figure S4B). Our findings that Netrin-1, but not Sema3A, induces the translation of mRNAs bound to DCC point towards a model where receptor-specific mRNA interactomes act as a hub for rapid cue-specific selective translation.

We next examined the effect of EphrinA1 co-stimulation on the Netrin-1-induced selective translation up-regulation of β-catenin, hnRNPH1 and RPS14/uS11. Quantification of immunofluorescence showed that EphrinA1 stimulation alone does not affect β-catenin and RPS14/uS11 protein levels (Figure 4D-G; S4C) and decreases hnRNPH1 protein level in axonal growth cones (Figure 4F-G). Co-stimulation with Netrin-1 and EphrinA1 blocks the Netrin-1-induced increase of all three proteins (Figure 4D-G; S4C). Together, the results show that integration of the EphrinA1 and Netrin-1 signals inhibits the Netrin-1-induced selective translation, possibly by inhibiting DCC-ribosome dissociation (Figure 3G).

### Ribosomes at the plasma membrane in axonal growth cones

Finally, we asked whether we could detect ribosomes in close proximity to the plasma membrane in axonal growth cones. We performed electron microscopy on axonal growth cones, and we observed an abundance of ribosomes in growth cones (Figure 5A). Strikingly, ribosomes could be seen aligned in rows underneath the plasma membrane (Figure 5A), particularly in the growth cone sections closest to the culture surface. These ribosomes were often single, rather than in clusters, indicative of and consistent with monosomes binding to the intracellular portions of transmembrane receptors such as DCC or Nrp-1.

**Figure 5.**
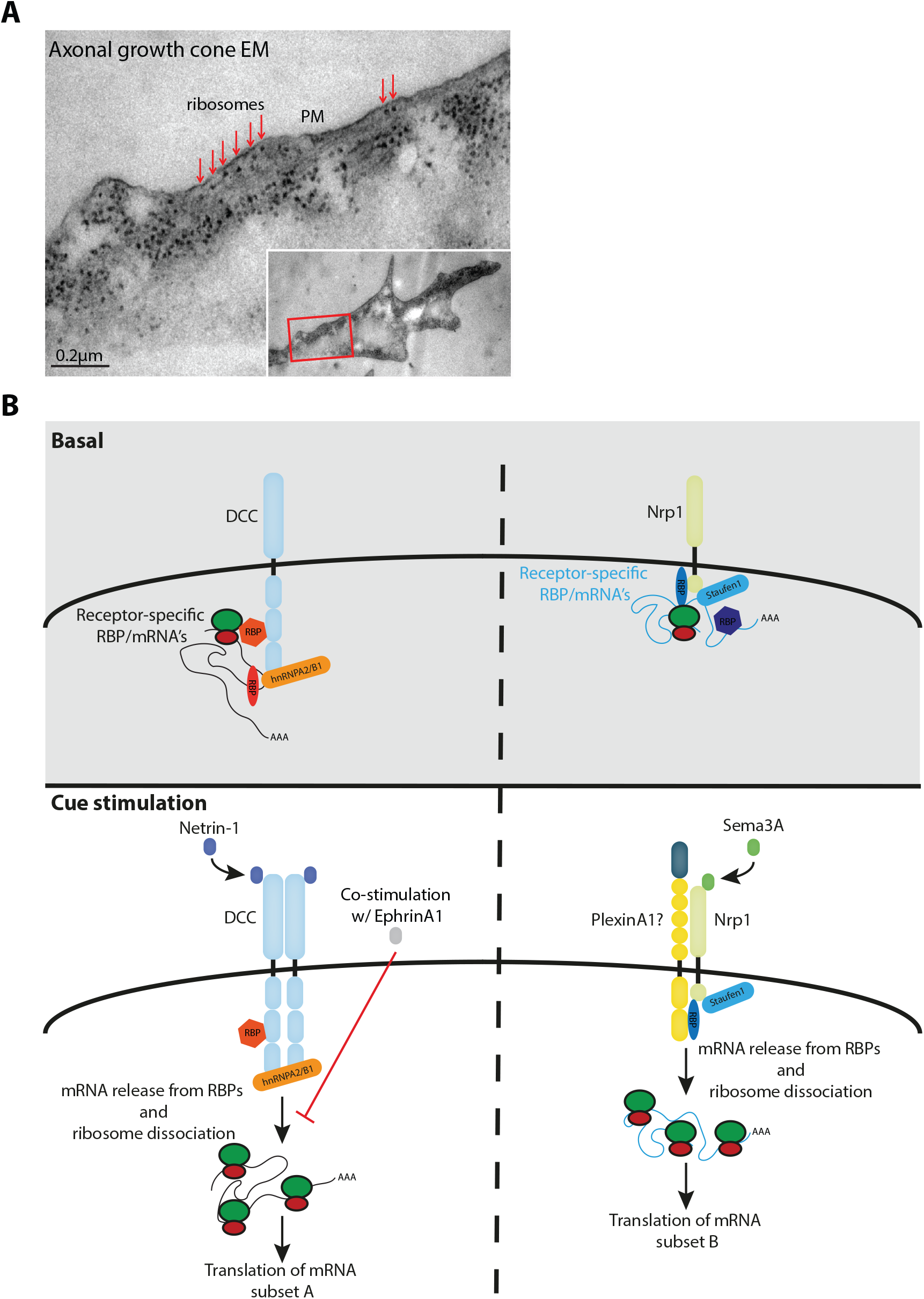
(A) EM image of axonal growth cone showing ribosomes aligned in a row (red arrows) under plasma membrane (PM). Inset shows the growth cone at lower magnification; the red-box indicates the area shown in higher magnification. The section glances through the extreme surface of growth cone, where it attaches to the culture dish, giving rise to areas that lack subcellular structure. (B) Model diagram depicting the proposed interactions between receptors, RBPs, mRNAs and ribosomes under basal and cue stimulation conditions.

## Discussion

We provide evidence for a receptor-ribosome coupled mechanism by which extrinsic cues cause rapid changes in the local proteome. In support of this model, we show that multiple guidance cue receptors interact with ribosomes, that the interaction between receptors and ribosomes depends on mRNA and rapidly decreases within 2 min of cue stimulation. Moreover, we show that receptors bind to distinct subsets of RBPs and mRNAs, and that cue stimulation induces the selective axonal translation of several receptor-specific mRNAs. Finally, we show that the integration of multiple cues can alter receptor-ribosome dissociation and selective translation.

Based on the candidate receptors tested here, we suggest that whether or not a particular receptor shows receptor-ribosome coupling is related to whether or not the receptors regulate local translation upon ligand binding. Future studies are needed to determine whether receptor-ribosome coupling is restricted to axon guidance receptors and neurons. Interestingly, a previous study has reported the association of a chemokine receptor, CXCR4, with eukaryotic initiation factor 2B (eIF2B), which decreases upon ligand binding (Palmesino et al., 2016). In addition, several adrenergic receptor subtypes have been reported to associate with eIF2B at the plasma membrane (Klein et al., 1997). This raises the intriguing possibility that coupling of translational machinery with receptors extends to other cell types and is a widespread mechanism to rapidly transduce local translation downstream of extracellular signals.

Label-free quantification of ribosomal proteins reveals possible differences in RP composition between DCC and Nrp1-bound ribosomes. Interestingly, both candidate RPs identified here, RPL4/uL4 and RPL39/eL39, are positioned near the peptide exit tunnel (Zhang et al., 2013) and could thereby influence translation rates (Wilson and Beckmann, 2011). Nevertheless, it should be noted that our current approach to analyse RP composition has several limitations which warrant caution. Firstly, label-free quantification is less accurate than quantification based on TMT-labels or single reaction monitoring. Secondly, these experiments did not include a ribosome isolation step, which raises the possibility that free ribosomal proteins bound to receptors contribute to our quantification. Therefore, future studies will be needed to investigate whether specific receptors bind heterogeneous ribosomes and whether these ribosomes are specialized to translate specific mRNA subsets. Ribosome heterogeneity can also be achieved by variation in ribosomal RNA content, ribosomal protein post-translational modification or association of different proteins with the ribosome (Slavov et al., 2015, Genuth and Barna, 2018, Shi et al., 2017, Emmott et al., 2018, Kurylo et al., 2018, Imami et al., 2018, Simsek et al., 2017), therefore, it will be interesting to investigate if these differences are present in receptor-bound ribosomes.

Previous studies have shown that the RBP zipcode binding protein 1 can be phosphorylated upon cue stimulation, thereby regulating local translation in axons by possibly releasing the bound mRNAs (Huttelmaier et al., 2005, Sasaki et al., 2010, Lepelletier et al., 2017). DCC and Nrp1 each differentially bind to RBPs and mRNAs, providing a way to rapidly achieve cue-induced selective translation. We observed an enrichment of known RBP targets for RBPs detected specifically in DCC and Nrp1 pulldowns respectively, suggesting a role for RBPs in mediating the differential binding of mRNAs to receptors and their cue-induced selective translation. This hypothesis is supported by the enrichment of the RBP hnRNPA2B1 specifically in DCC but not Nrp1 pulldown, as hnRNPA2B1 has been reported to control the translation of β-catenin (Stockley et al., 2014), which is selectively translated in response to Netrin-1, but not Sema3A (Cagnetta et al., 2018), in accord with the interactome data reported here.

Our results point to a model in which different subsets of mRNAs interact via specific RBPs with either DCC or Nrp1, and are released, together with ribosomes, upon specific cue stimulation and thus become available for subsequent translation (Figure 5; model). It should be noted that, in addition to RBPs and mRNAs, several other molecules characterize the receptor-specific interactome. For example, eIF3d, an initiation factor previously shown to regulate specialized translation initiation, is significantly enriched specifically in Nrp1 IP, but not DCC IP, thus raising the interesting possibility that differential binding to initiation factors may contribute to cue-induced selective translation (Lee et al., 2016). Intriguingly, a recent study revealed that an untranslated mRNA can associate with and regulate the signalling of the TrkA receptor in axons via its axon-enriched long 3’UTR (Crerar et al., 2019). It will be interesting to investigate whether any of the DCC and Nrp1 targets identified in our study also play a structural role, for example by regulating the receptor-ribosome association and/or the downstream signalling and local translation.

During axon guidance and branching, axons can encounter a combination of extracellular signals and ample evidence shows that the integration of multiple cues results in different outcomes than those of each single cue (Dudanova and Klein, 2013, Morales and Kania, 2017). Here, we tested the effect of cue integration on receptor-ribosome coupling and found that EphrinA1 blocks Netrin-1 induced ribosome dissociation from DCC. In addition, EphrinA1 blocks the Netrin-1-induced selective increase in translation of several mRNAs. The mechanism by which EphrinA1 affects the coupling of DCC to ribosomes is unknown. One possibility is that, upon co-stimulation of EphrinA1 and Netrin-1, the DCC and Eph receptors may form a complex, thereby altering the receptor structure and association to ribosomes, which could be consistent with a previous study showing a ligand-dependent interaction between the receptors Unc5 and EphB2 (Poliak et al., 2015).

In conclusion, our findings show that coupling of the translational machinery to guidance cue receptors at the plasma membrane of growth cones is a mechanism to rapidly and selectively control the cue-induced regulation of the local proteome and suggest that this is a general principle that can be applied to neural membrane receptors more broadly.

---

**Figure S1.**
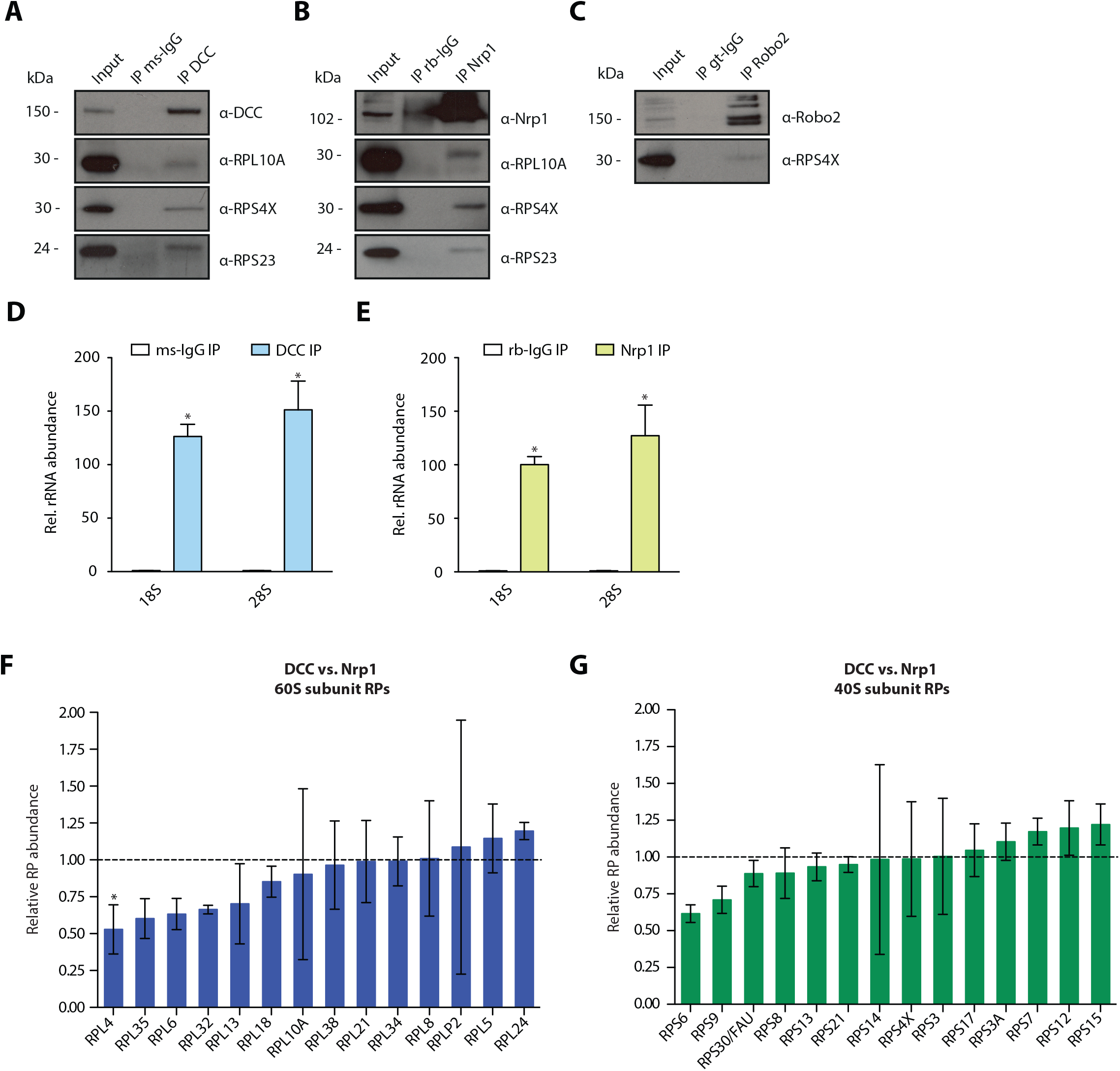
Multiple guidance cue receptors interact with ribosome in SH-SY5Y cells. (A-C) Western blot validation of RP co-immunoprecipitation with DCC, Nrp1 and Robo2 in SH-SY5Y cells. (D-E) Relative 18S and 28S ribosomal RNA abundance after control (IgG) pulldowns or receptor pulldowns shows enrichment of rRNA in DCC and Nrp1 IPs in SH-SY5Y cells (unpaired two-tailed t-test; three biological replicates). (F) iBAQ-based relative quantification of 60S or (G) 40S subunit RPs between DCC and Nrp1 (unpaired two-tailed t-test; three biological replicates). Error bars indicate standard deviation. *p<0.05.

**Figure S2.**
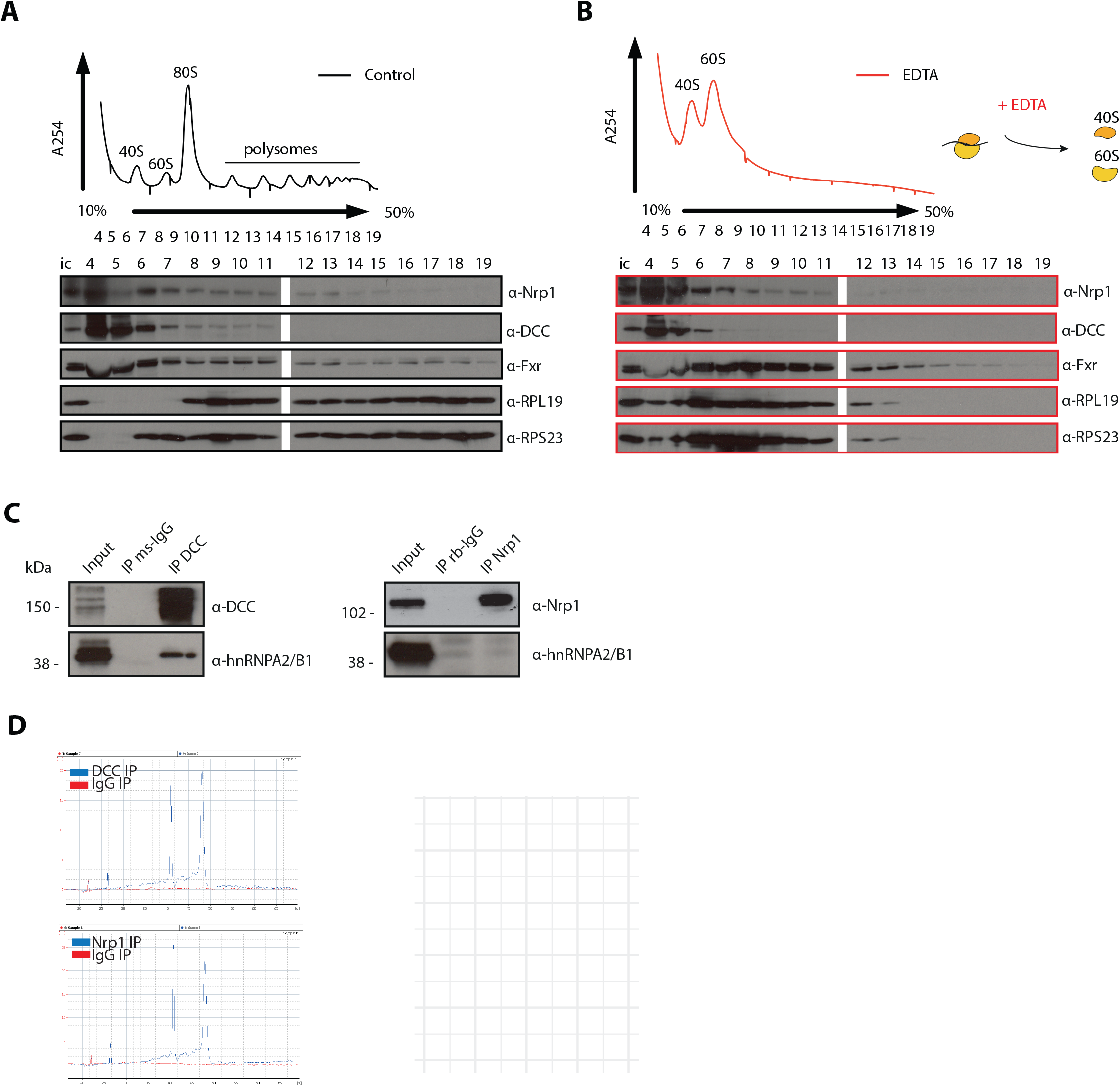
(A) Control and (B) EDTA treated polysome fractions and Western blot showing the distribution of DCC and Nrp1 across fractions. (C) Western blot showing hnRNPA2/B1 is co-immunoprecipitating with DCC but not with Nrp1. (D) Bioanalyzer gel analysis of RNA. (E) Gene ontology enrichment plot of mRNAs after DCC or (F) Nrp1 pulldowns.

**Figure S3.**
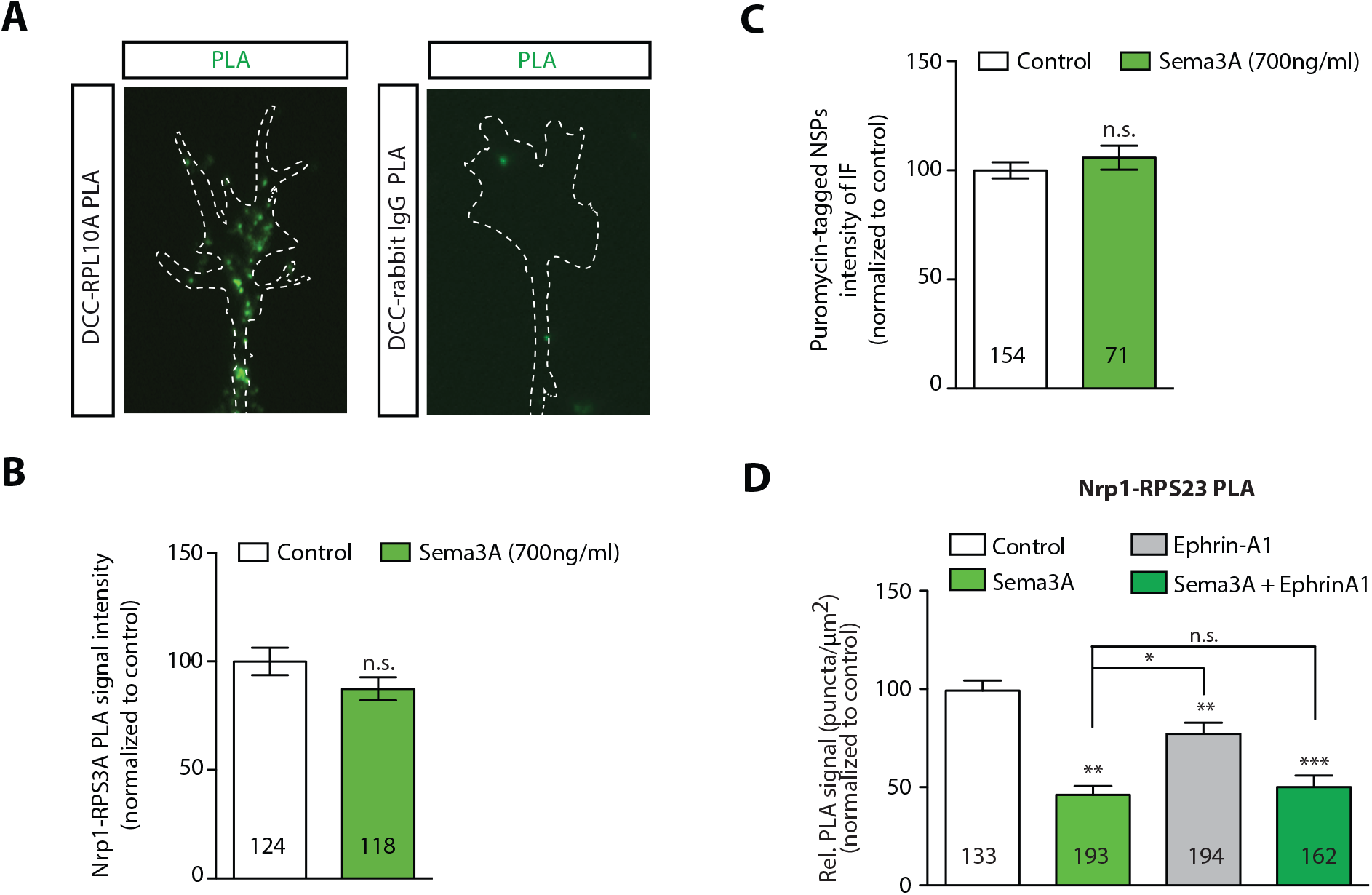
(A) PLA images showing DCC and RPL10A/uL1 are in close proximity in axonal growth cones, whereas DCC and IgG control generates little to no PLA signal. Scale bars, 5 μm. (B) Sema3A stimulation at protein-synthesis independent concentration does not decrease PLA signal between Nrp1 and RPS3A/eS1 (Mann-Whitney test; error bars indicate SEM; p = 0.2555). or (C) puromycin levels in axonal growth cones (Mann-Whitney test; error bars indicate SEM; p = 0.2487). (D) Relative PLA quantification of Nrp1 and RPS23/uS12 compared to control after Sema3A, EphrinA1, or co-stimulation with Sema3A and EphrinA1 (one-way ANOVA with Bonferroni’s multiple comparisons test; Error bars indicate SEM; *p=0.032078; **p<0.018577; *\*\*\**p<0.001). For all PLA and QIF experiments, numbers in bars indicate amount of growth cones quantified collected from at least three independent experiments.

**Figure S4.**
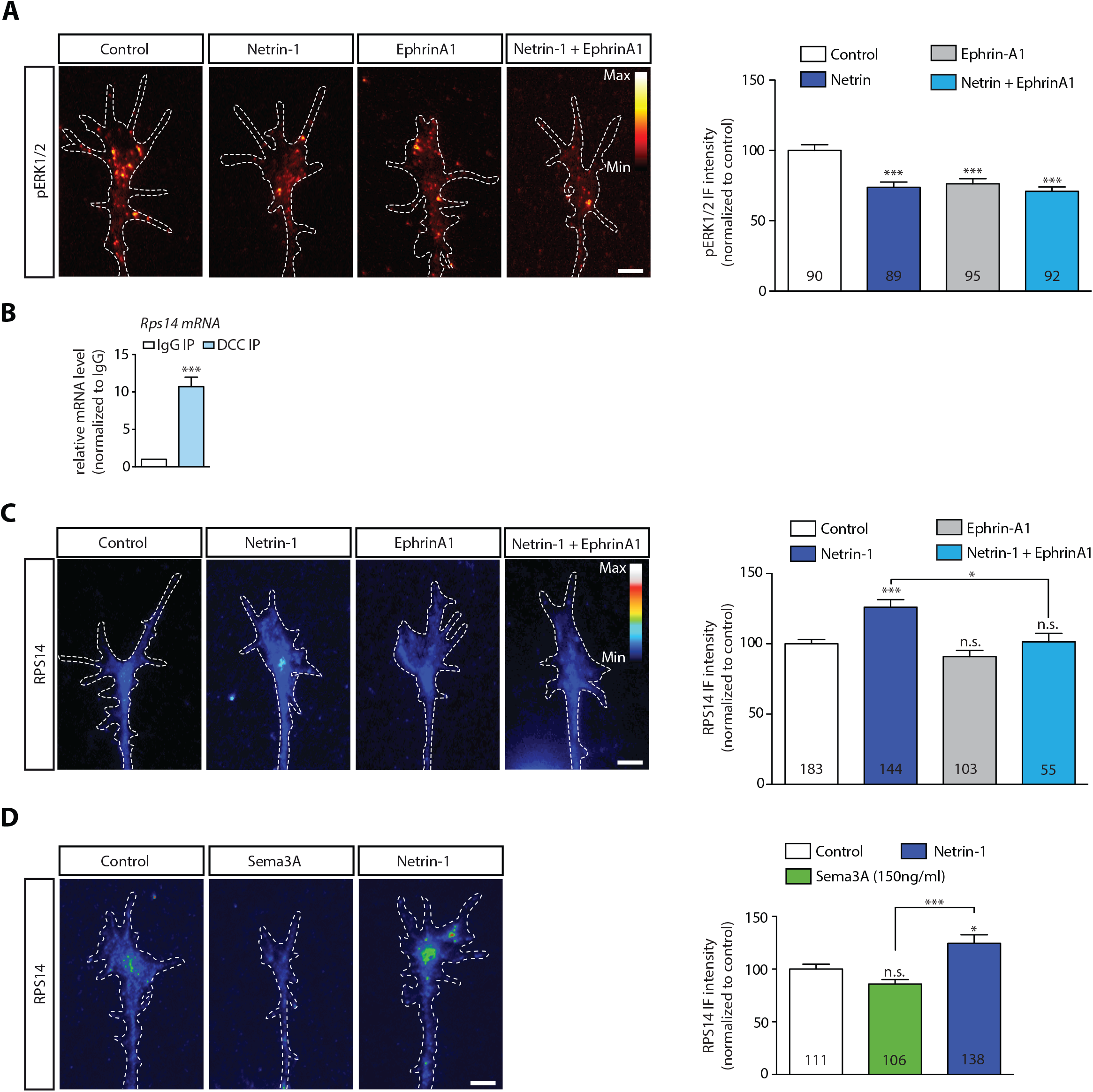
(A) Netrin-1 and EphrinA1 both decrease pERK1/2 levels in axonal growth cones. RPS14/uS11 QIF relative to control after Netrin-1, EphrinA1, or co-stimulation. (B) Relative mRNA quantification after DCC IP of *rps14* mRNA (unpaired t-test with Welch’s corrections on dCT values; three biological replicates; error bars indicate SEM; ***p = 0.0003). (C) Rps14 QIF relative to control after Netrin-1, EphrinA1 or Netrin-1 and EphrinA1 co-stimulation (one-way ANOVA with Bonferroni’s multiple comparisons test; error bars indicate SEM; ***p<0.0001; *p=0.026544). (D) Rps14 QIF relative to control after Netrin-1 or Sema3A stimulation (one-way ANOVA with Bonferroni’s multiple comparisons test; error bars indicate SEM; ***p<0.0001; *p<0.05). Scale bars, 5 μm. For all QIF experiments the numbers in bars indicate amount of growth cones quantified collected from at least three independent experiments.

## Materials and methods

### Embryos

*Xenopus laevis* embryos were fertilized *in vitro* and raised in 0.1x Modified Barth’s Saline (8.8mM NaCl, 0.1 mM KCl, 0.24mM NaHCO_3_, 0.1 mM HEPES, 82µM MgSO_4_, 33µM Ca(NO_3_)_2_, 41µM CaCl_2_) at 14-20°C and staged according to the tables of Nieuwkoop and Faber (Nieuwkoop and Faber, 1994). All animal experiments were approved by the University of Cambridge Ethical Review Committee in compliance with the University of Cambridge Animal Welfare Policy.

### Primary *Xenopus* retinal cultures

Eye primordia were dissected from Tricaine Methanesulfonate (MS222) (Sigma-Aldrich) anesthetized embryos at stage 35/36 (or stage 32 for EM) and cultured on 10µg/ml poly-L-lysine-(Sigma-Aldrich) and 10µg/ml laminin-(Sigma-Aldrich) coated dishes in 60% L-15 medium (Gibco) at 20°C for 24h before performing immunohistochemistry or proximity ligation assay, or for 48h before the puromycilation assay.

### Immunoprecipitation

SH-SY5Y cells or Xenopus brains and eyes dissected from stage 40/41 embryos were lysed in lysis buffer (20mM Tris-HCl, pH 7.4, 150mM NaCl, 10mM MgCl2 and 10% glycerol supplemented with 100µg/ml cycloheximide (Sigma-Aldrich), EDTA-free protease inhibitors (Roche, 11873580001), phosphatase inhibitors (Thermo Fisher Scientific, A32957) and SuperRNAse In RNAse inhibitor (Ambion, AM2696)). Tissues or cells were lysed for 30 minutes at 4C and centrifuged for 5 minutes at 800g at 4°C to remove unlysed cells and nuclei and then 15 minutes at 16000g at 4°C. The resulting supernatant was incubated with magnetic Dynabeads pre-coupled with antibodies using the Dynabeads antibody coupling kit (Thermo Fisher Scientific, 14311D) for 1.5 hours at 4°C on a rotor. The following antibodies were used: mouse-anti-DCC (BD Biosciences, 554223); rabbit-anti-Nrp1 (Abcam, ab81321); goat-anti-Robo2 (R&D systems, AF3147); mouse-anti-EphB2 (Santa Cruz, sc130068) or an isotype control: rabbit IgG (Abcam, ab37415); mouse IgG1 (R&D systems, 11); mouse IgG2b (R&D systems, MAB004); goat IgG (R&D systems, AB-108-C). Beads were then washed 3 times in lysis buffer and processed for protein or RNA isolation. For EDTA treatment, beads were washed with EDTA wash buffer (20mM Tris-HCl, pH 7.4, 150mM NaCl, 25mM EDTA and 10% glycerol supplemented with EDTA-free protease inhibitors (Roche, 11873580001), phosphatase inhibitors (Thermo Fisher Scientific, A32957) for 3 times before elution. For RNaseA/T1 treatment, beads were washed three times for 3 minutes at RT with RNaseA/T1 wash buffer (20mM Tris-HCl, pH 7.4, 150mM NaCl, 10mM MgCl2 and 10% glycerol supplemented with 100µg/ml cycloheximide (Sigma-Aldrich), EDTA-free protease inhibitors (Roche, 11873580001), phosphatase inhibitors (Thermo Fisher Scientific, A32957), 10µg/µl RNase A (Ambion, EN0531) and 250U RNase T1 (Ambion, EN0541). The concentrations of RNaseA and RNase T1 used here were previously determined to selectively affect polysomes but not monosomes using polysome fractionation analysis (Shigeoka et al., 2018).

For protein isolation, 1x NuPAGE LDS sample buffer (Thermo Fisher Scientific, NP0008) was added to the beads, incubated for 5 minutes at 95°C and the final protein eluate was collected after magnetic separation of the beads. For RNA isolation, RLT buffer was added to the beads, vortexed for 2 minutes and then separated from the beads on a magnetic stand.

### Polysome fractionation

For density gradient fractionation, lysate was layered on a sucrose gradient (10-50%) in PLB buffer (20mM Tris-HCl, pH 7.4, 150mM NaCl, 10mM MgCl2, 100µg/ml cycloheximide (Sigma-Aldrich), 0.5mM DTT) and ultracentrifugation was performed using a Beckman SW-40Ti rotor and Beckman Optima L-100 XP ultracentrifuge, with a speed of 35,000 rpm at 4°C for 160 min. Fractionations and UV absorbance profiling were carried out using Density Gradient Fractionation System (Teledyne ISCO). Proteins were precipitated from each fraction using methanol-chloroform precipitation and pellets were resuspended in 1x NuPAGE LDS sample buffer and used for Western blotting as described below.

### Western blot

Proteins were resolved by SDS-PAGE on NuPage 4-12% Bis-Tris gels (Invitrogen, NP0321) and transferred to nitrocellulose membrane (Bio-Rad). The blots were blocked in 5% milk in TBST-T for 60 minutes at RT and then incubated with primary antibodies in 5% milk in TBS-T overnight at 4°C. After washing 3 times with TBS-T the blots were incubated with HRP-conjugated secondary antibodies (goat-anti-mouse HRP (Abcam, ab6789); goat-anti-rabbit HRP (Abcam, ab6721) for 1 hour at RT, washed again for 3 times in TBS-T, followed by ECL-based detection (Pierce ECL plus, Thermo Scientific, 32123). The following primary antibodies were used for Western blot analysis: mouse-anti-DCC (BD Biosciences, 554223), rabbit-anti-neuropilin-1 (Abcam, ab81321), goat-anti-Robo2 (R&D systems, AF3147), mouse-anti-EphB2 (Santa Cruz, sc130068), mouse anti-Rpl19/eL19 (Abcam, ab58328), mouse anti-RPS23/uS12 (Abcam, ab57644), rabbit anti-RPS4X/eS4 (Proteintech, 14799-1-AP), rabbit-anti RPL10A/uL1 (Proteintech, 16681-1-AP), rabbit-anti Rps26 (Proteintech, 14909-1-AP), mouse-anti-Rps3A (Abcam, ab194670), mouse-anti-FxR (gift from dr. Khandjian), rabbit-anti-hnRNPA2/B1 (Abcam, ab31645).

### Quantitative RT-PCR

RNA was isolated from eluted samples using the RNeasy Mini kit (Qiagen, 74104) and reverse transcribed into cDNA using random hexamers and the SuperScript III First-Strand Synthesis System (Thermo Fisher Scientific, 18080051). The cDNA was used to prepare triplicate reactions for qRT-PCR according to manufacturer’s instructions (QuantiTect SYBR Green PCR kit, Qiagen, 204143), plates were centrifuged shortly and run on a LightCycler 480 (Roche) using the following PCR conditions: denaturation for 15s at 94°C; annealing for 30s at 60°C; extension for 30s at 72°C. The levels for each condition were corrected with their own input. The following primers were used for qPCR: *Xenopus 18S rRNA*, 5’-GTAACCCGCTGAACCCCGTT-3’ and 5’-CCATCCAATCGGTAGTAGCG-3’; *Xenopus 28S rRNA*, 5’-CTGTCAAACCGTAACGCAGG-3’ and 5’-CTGACTTAGAGGCGTTCAGTCA-3’. *human 18S rRNA*, 5’-GTAACCCGTTGAACCCCATT-3’ and 5’-CCATCCAATCGGTAGTAGCG-3’; *human 28S rRNA*, 5’-AACGGCGGGAGTAACTATGA-3’ and 5’-TAGGGACAGTGGGAATCTCG-3’. Xenopus Ctnnb1 mRNA, 5’-GACCACAAGTCGGGTGCTTA-3’ and 5’-CCAGACGTTGGCTTGAGTCT-3’; Xenopus hnrnph1 mRNA, 5’-GGTTGGAAAATCGTGCCAAATG-3’ and 5’-GCCTTTTCAGCTATTTCCTGTGAAG-3’; Xenopus rps14 mRNA, 5’-GTGACTGACCTGTCTGGCAA-3’ and 5’-GCAACATCTTGTGCAGCCAA-3’.

### Proximity ligation assay

This experiment was carried out according to the manufacturer’s protocol (Sigma-Aldrich, Duolink Biosciences) using Duolink In Situ Detection reagents (Sigma-Aldrich, DUO90214 or DUO92008). After 24h, cultures were fixed in 2% formaldehyde/7.5% sucrose in PBS for 20 minutes at 20°C, washed 3 times in PBS with 0.001% Triton-X-100, permeabilized for 5 minutes in 0.1% Triton-X-100 in PBS, washed three times in PBS with 0.001% Triton-X-100, blocked with 5% heat-inactivated goat serum in PBS for 45 minutes at RT and subsequently incubated with primary antibodies overnight at 4°C. Primary antibodies were diluted at 1:100 for mouse anti-DCC (BD Biosciences, 554223), 1:100 mouse-anti-EphB2 (Thermo Fisher Scientific, 137-1700) 1:100 for rabbit anti-RPL5/uL18 (Proteintech, 15430-1-AP), 1:100 rabbit anti-RPS4X/eS4 (Proteintech, 14799-1-AP), 1:100 rabbit-anti RPL10A/uL1 (Proteintech, 16681-1-AP), 1:100 for rabbit anti-neuropilin-1 (Abcam, ab81321), 1:100 mouse anti-RPS3A/eS1 (ab194670, Abcam),1:100 mouse-anti-RPS23/uS12 (ab57644, Abcam), rabbit-IgG isotype control (ab37415, Abcam), mouse IgG1 isotype control (11711, R&D Systems). After primary antibody incubation, dishes were washed twice for 5 minutes with 0.002% Triton X-100 in PBS and incubated with anti-rabbit-PLUS (Sigma-Aldrich, DUO92002) and anti-mouse-MINUS (Sigma-Aldrich, DUO92004) PLA probes for 1 hour at 37°C, with ligase for 30 minutes at 37°C and with the polymerase mix with red fluorescence for 100-140 min at 37°C. The samples were subsequently mounted with the mounting medium (DUO82040, Duolink) and imaged using a Nikon Eclipse TE2000-U inverted microscope equipped with an EMCCD camera. The number of discrete fluorescent puncta from randomly selected isolated growth cones were counted using Volocity software (Perkin Elmer).

### Immunocytochemistry

After 24 hours, Xenopus retinal cultures were fixed in 2% formaldehyde/7,5% sucrose in PBS for 20 min at 20°C. For the puromycilation assay, 48h old cultures, eyes were manually removed and axons were treated with 10µg/ml puromycin for 10 minutes at RT before fixation. The fixed cultures were then washed 3 times in PBS with 0.001% Triton-X-100, permeabilized for 5 min at RT in 0.1% Triton-X-100 in PBS, washed again for three time in PBS with 0.001% Triton-x-100 and blocked with 5% heat-inactivated goat serum in PBS for 45 min at 20°C. Primary antibodies were incubated overnight at 4°C, followed by Alexa Fluor-conjugated secondary antibodies for 60 min at 20°C in the dark. Cultures were mounted in FluorSave (Calbiochem, 345789). Primary antibodies were used at the following dilutions: 1:100 for mouse anti-DCC (BD Biosciences, 554223), 1:100 for rabbit anti-neuropilin-1 (Abcam, ab81321), 1:100 for rabbit anti-RPL5/uL18 (15430-1-AP, Proteintech), 1:100 mouse anti-RPS3A/eS1 (ab194670, Abcam), 1:200 mouse-anti-puromycin-AlexaFluor-488 (Millipore, MABE343-AF488), 1:500 rabbit-anti-β-Catenin (Sigma-Aldrich, C2206), 1:500 rabbit-anti-hnRNPH1 (Abcam, ab154894), rabbit-anti-RPS14/uS11 (Abcam,), 1:250 rabbit-anti-pERK1/2 (Cell Signaling, 9101). Secondary antibodies were diluted at: 1:1000 goat anti-rabbit Alexa Fluor 568 (Abcam, ab150077), 1:1000 goat anti-mouse Alexa Fluor 568 (Abcam, ab150117).

### Expansion microscopy

For expansion microscopy, RGCs explant cultures were immunostained with primary and secondary antibodies as described above, followed by applying the expansion protocol for cultured cells (Chen et al., 2015). Briefly, cultures were incubated in 0.25% glutaraldehyde in PBS for 20 min at RT and then washed with PBS three times, before adding monomer solution (2M NaCl, 8.625% (w/w) sodium acrylate, 2.5% (w/w) acrylamide, 0.1% (w/w) N,N‘-methylenebisacrylamide in PBS) for 2 min at RT. Subsequently, monomer solution was mixed with 0.2% ammonium persulfate (APS) and 0.2% Tetramethylethylendiamin (TEMED) and added to the samples. Gelation of the polymer occurred at 37°C for 30 min, followed by digestion of the samples with digestion buffer (40mM Tris (pH 8), 1mM EDTA, 0.5% Triton-X-100, 0.8M guanidine NaCl, 8U/ml Proteinase K in water) and incubated at 37°C for 1h. To expand the samples, digestion buffer was removed and gels were placed in water for several hours during which water was replaced every 30 min. Once gels detached from the glass dish, they were transferred to a bigger dish to allow expansion. For imaging, expanded gels were cut in pieces and transferred to poly-L-lysine coated glass bottom dishes. Imaging was performed using a 60x/1.3 NA silicone oil objective lens on a Perkin Elmer Spinning Disk UltraVIEW ERS, Olympus IX81 inverted microscope and the Volocity software.

### Quantification of Immunofluorescence

For the quantification of fluorescence intensity, isolated growth cones were randomly selected with phase optics. For each experiment, the images were captured on the same day using the same gain and exposure settings and pixel saturation was avoided. Using Volocity software (Perkin Elmer), a region of interest (ROI) was defined by tracing the outline of each single growth cone using the phase image and the mean pixel intensity per unit area was measured in the fluorescent channel. The background fluorescence was measured in a ROI close to the growth cone that was free of debree or other axons and this was substracted from the mean fluorescence value of the growth cone.

### Mass-spectrometry

1D gel bands were transferred into a 96-well PCR plate. The bands were cut into 1mm2 pieces, destained, reduced (DTT) and alkylated (iodoacetamide) and subjected to enzymatic digestion with chymotrypsin overnight at 37°C. After digestion, the supernatant was pipetted into a sample vial and loaded onto an autosampler for automated LC-MS/MS analysis. All LC-MS/MS experiments were performed using a Dionex Ultimate 3000 RSLC nanoUPLC (Thermo Fisher Scientific Inc, Waltham, MA, USA) system and a Q Exactive Orbitrap mass spectrometer (Thermo Fisher Scientific Inc, Waltham, MA, USA). Separation of peptides was performed by reverse-phase chromatography at a flow rate of 300nL/min and a Thermo Scientific reverse-phase nano Easy-spray column (Thermo Scientific PepMap C18, 2μm particle size, 100A pore size, 75μm i.d. x 50cm length). Peptides were loaded onto a pre-column (Thermo Scientific PepMap 100 C18, 5μm particle size, 100A pore size, 300μm i.d. x 5mm length) from the Ultimate 3000 autosampler with 0.1% formic acid for 3 minutes at a flow rate of 10μL/min. After this period, the column valve was switched to allow elution of peptides from the pre-column onto the analytical column. Solvent A was water + 0.1% formic acid and solvent B was 80% acetonitrile, 20% water + 0.1% formic acid. The linear gradient employed was 2-40% B in 30 minutes. The LC eluant was sprayed into the mass spectrometer by means of an Easy-Spray source (Thermo Fisher Scientific Inc.). All m/z values of eluting ions were measured in an Orbitrap mass analyzer, set at a resolution of 70000 and was scanned between m/z 380-1500. Data-dependent scans (Top 20) were employed to automatically isolate and generate fragment ions by higher energy collisional dissociation (HCD, NCE:25%) in the HCD collision cell and measurement of the resulting fragment ions was performed in the Orbitrap analyser, set at a resolution of 17500. Singly charged ions and ions with unassigned charge states were excluded from being selected for MS/MS and a dynamic exclusion window of 20 seconds was employed.

Raw data were processed using Maxquant (version 1.6.1.0) (Cox and Mann, 2008) with default settings. MS/MS spectra were searched against the *X. laevis* protein sequences from Xenbase (xlaevisProtein.fasta). Enzyme specificity was set to trypsin/P, allowing a maximum of two missed cleavages. The minimal peptide length allowed was set to seven amino acids. Global false discovery rates for peptide and protein identification were set to 1%. The match-between runs option was enabled.

### Label-free quantification (LFQ/iBAQ) analysis of proteomics data

To identify significant interactors, t-test-based statistics were applied on label-free quantification (LFQ) intensity values were performed using Perseus software. Briefly, LFQ intensity values were logarithmized (log2) and missing values were imputed based on the normal distribution (width = 0.3, shift = 1.8). Significant interactors of DCC or Nrp1 pulldowns compared to IgG pulldowns were determined using a two-tailed t-test with correction for multiple testing using a permutation-based false discovery rate (FDR) method. To determine the stoichiometry of ribosomal proteins in 40S and 60S subunits, we compared the relative abundance of all detected ribosomal proteins as measured by iBAQ intensities. The obtained relative abundances were normalized to the mean of all iBAQ intensities for either 40S or 60S ribosomal proteins in each sample. These normalized relative abundances were then compared between DCC and Nrp1 and tested for significance using a t-test.

### RNA-sequencing

RNA was isolated from immunoprecipitated samples from SH-SY5Y cells as described above using RLT buffer (Qiagen) containing β-mercaptoethanol and the RNeasy Mini kit (Qiagen) followed by in-column DNase I treatment to remove genomic DNA contamination. RNA quality was analysed using Agilent RNA 6000 Pico kit and reagents (Agilent, 5067-1514,1535,1513) on a Agilent 2100 Bioanalyzer (Agilent). cDNA was then amplified using a method developed for single cell transcriptomics (Tang et al., 2009) with minor modifications (Shigeoka et al., 2016). The cDNA library preparation was performed using a KAPA Hyperprep kit (Roche) and cDNA libraries were subjected to a RNA-sequencing run on a Next-seq 500 instrument (Illumina) using the 150 cycles high output kit (Illumina).

### Bioinformatic analysis of RNA-sequencing data

The sequence reads were mapped using HISAT 2 version 2.1.0, and FPKM values were estimated using Cufflinks version 2.1.1. Read counts for each gene were determined using HTSeq version 0.11.0. Differential expression analysis was performed using edgeR in R version 3.5.0 (FDR < 0.05). The GO enrichment analysis was performed using topGO version 2.32.0. The mRNA targets of RBPs were obtained from previously published studies as listed in the main text.

### Electron microscopy of axonal growth cones

Cultured neurons were fixed at 37°C for 45 min in 2.5% glutaraldehyde, sodium cacodylate buffer 0.1M pH7.4 containing 2mM CaCl2 and 2mM MgCl2. Samples were post-fixed for 15min at RT in 1% osmium and embedded in epoxy resin. Ultrathin sections were imaged with a ZEISS EM 912 microscope. Ribosomes were identified based on size and shape.

## Author Contributions

M.K. and C.E.H. designed the experiments. M.K. wrote the manuscript with support from R.C., T.S., J.F., and C.E.H. M.K performed all biochemical experiments. M.K., R.C., S.Z and M.M. carried out *in vitro* retinal cultures, immunohistochemistry and PLA. L.W. and M.K. performed expansion microscopy. M.K. performed RIP-seq experiments. T.S and M.K. performed bioinformatics analysis. A.B. performed electron microscopy. C.E.H., C.F.K., and M.K. supervised the project.

## Acknowledgements

We thank William A. Harris for valuable input on the manuscript. We thank Nicola Lawrence, Caia Duncan (Juan Mata lab, Unversity of Cambridge), Katrin Mooslehner and Asha Dwivedy for technical assistance and Fabrice Richard (PiCSL-FBI core facility, IBDM, CNRS, Aix-Marseille University; member of the France-BioImaging national research infrastructure (ANR-10-INBS-04)) for assistance with EM experiments. This work was supported by the Netherlands Organization for Scientific Research (NWO Rubicon 019.161LW.033) (M.K.), Wellcome Trust Grants (085314/Z/08/Z and 203249/Z/16/Z) and European Research Council Advanced Grant (322817) (C.E.H.).

## Competing interests

The authors declare no competing interests.

